# Genomic and morphological characterisation of a novel iridovirus, bivalve iridovirus 1 (BiIV1), infecting the common cockle (*Cerastoderma edule*)

**DOI:** 10.1101/2025.03.05.641634

**Authors:** Chantelle Hooper, Anna M. Tidy, Ron Jessop, Kelly S. Bateman, Matthew J. Green, Stuart H. Ross, Georgia M. Ward, Richard Hazelgrove, Jasmine E. Hunt, Megan Parker, David Bass

**Affiliations:** International Centre of Excellence for Aquatic Animal Health, The Centre for Environment, Fisheries and Aquaculture Science, Weymouth, DT4 8UB, UK; Sustainable Aquaculture Futures, Biosciences, Faculty of Health and Life Sciences, University of Exeter, Exeter, EX4 4QD, UK; Eastern Inshore Fisheries & Conservation Authority (EIFCA), North Lynn Business Village, Bergen Way, North Lynn Industrial Estate, King’s Lynn, PE30 2JG, UK; School of Biosciences and Medicine, University of Surrey, Guildford, Surrey, GU2 7XH, UK

**Keywords:** iridovirus, cockle, bivalve, emerging disease, aquatic animal health

## Abstract

High rates of mortality of the common cockle, *Cerastoderma edule*, have occurred in the Wash Estuary, UK since 2008. A previous study linked the mortalities to a novel genotype of *Marteilia cocosarum*, with a strong correlation between cockle moribundity and the presence of *M. cocosarum*. Here, we characterise a novel iridovirus, identified by chance during metagenomic sequencing of a gradient purification of *Marteilia* cells, with presence also correlated to cockle moribundity. The novel 179,695 bp iridovirus, bivalve iridovirus 1 (BiIV1), encodes 193 predicted open reading frames and has a 41% G+C content. BiIV1 clusters together with other aquatic invertebrate iridoviruses in phylogenetic analyses and has a similar genome size to other invertebrate iridoviruses. Comparative analysis revealed that BiIV1 has lost three genes that were previously thought to be common amongst all iridoviruses but has also gained genes, potentially from horizontal transfer from its bivalve mollusc host(s). Electron microscopy showed 158 nm icosahedral virions present in the haemocytes of cockles, typical of those observed in host tissues infected with viruses of the family *Iridoviridae*. Prevalence of BiIV1 in moribund cockles was higher than that in apparently healthy cockles at most sites in the Wash Estuary, with up to 100% PCR prevalence in moribund cockles. Our findings provide the first genome for a bivalve-infecting iridovirus and identify a second bivalve-associated iridovirus in publicly available genomic datasets, adding to the knowledge of invertebrate iridovirus genomics and diversity.

## 2 Introduction

The common cockle, *Cerastoderma edule*, is a commercially important bivalve in Europe, with 25,000 tonnes produced in 2021 (FAO 2022). The United Kingdom is a large producer of cockles, producing a total of 7,500 tonnes in 2021, with cockles harvested from managed and wild fisheries. Cockles are known to be infected with many eukaryotic parasites including trematode and nematode worms, copepods and decapods, microsporidia, Alveolata, and Rhizaria (de Montaudouin et al. 2021). Despite the large number of eukaryotic parasites known to infect cockles, the presence of them in populations is typically low and detrimental effects due their presence is unlikely (de Montaudouin et al. 2021). However, infection with a rhizarian parasite, *Marteilia cochillia*, has been the biggest pathogen-associated threat to cockle production, with high presence and pathogenicity in populations in Spain causing mass mortalities (Carrasco et al. 2013).

*Marteilia cochillia* is a protistan parasite in the order Paramyxida (Rhizaria). *M. cochillia* was characterised in *C. edule* from Alfacs Bay (Mediterranean coast of Spain) in 2013 (Carrasco et al. 2013) but has been associated with mass mortalities in *C. edule* since 2008 (Carrasco et al. 2011). *M. cochillia* has since been determined as the cause of collapse of the *C. edule* fishery in Ría de Arousa (NW Spain), where mortalities reached 100% in 2012 (Villalba et al. 2014). Histological analysis of infected cockles showed heavy presence of the parasite in the digestive gland, with all tubules within a histology section frequently infected (Villalba et al. 2014). In 2017, a novel *Marteilia* parasite, *M. cocosarum*, was detected in Wales, UK (Skujina et al. 2022). Mortalities had been observed in the locations where *M. cocosarum* had been detected, but not to the extent seen in Spain, and could not be directly linked to infection with *Marteilia*. The tissue tropism of *M. cocosarum* was not typical of other *Marteilia* species and showed marked differences from that of infection with *M. cochillia*, with infection seen primarily in gill and mantle tissue and the connective tissue surrounding the digestive gland. Limited screens showed that *M. cocosarum* appeared to be geographically localised to cockle beds in Wales – Skujina et al. (2022) found *M. cocosarum* absent from four sites around the coast of England and four sites around the coast of Ireland.

Recently, *M. cocosarum* has been associated with *C. edule* mortalities in the Wash Estuary (Tidy et al. 2025). The Wash Estuary, located on the East coast of the UK is a fishery of many important aquatic species including cockles, mussels and brown shrimp (Hormbrey 2018). Mortalities of cockles have occurred in the Wash Estuary since 2008. Tidy et al. (2025) sampled moribund and healthy cockles over two years, finding a significantly higher presence of *M. cocosarum* in the moribund animals. Histopathology of moribund cockles showed that infection with *M. cocosarum* in cockles from the Wash had an identical phenotype to infections in Wales, but phylogenetic analysis of the ribosomal RNA array showed that *M. cocosarum* from the Wash Estuary was genetically distinct from *M. cocosarum* from Wales, therefore deeming it a new genotype: *M. cocosarum* WE. However, presence of *M. cocosarum* WE was not associated with all mortalities of cockles in the Wash.

Known bacterial and viral agents that infect cockles are relatively few compared to their eukaryotic parasites. The only bacterial species reported to be a threat to cockle production is *Vibrio aestuarianus*, also responsible for mortalities of other bivalve species, and has been identified as the causative agent of cockle summer mortality events in France (Garcia et al. 2021). Viral pathogens of cockles are rarely described, however an undescribed picornavirus-like infection has been associated with granulomatosis (Carballal et al. 2003), viral lesions have been observed in the digestive tubules of cockles infected with *M. cochillia* (Carrasco et al. 2011), and *C. edule* can become infected with oyster herpesvirus (OsHV-1) (Bookelaar et al. 2020).

In this study, we describe a novel iridovirus detected by chance during gradient purification of *Marteilia* parasites from moribund cockles infected with *M. cocosarum* WE from the Wash Estuary in 2022. Iridoviruses are double-stranded DNA (dsDNA), icosahedral viruses typically 120-200 nm in diameter. *Iridoviridae* comprises two subfamilies, *Alphairidovirinae* and *Betairidovirinae*, with the former infecting primarily ectothermic vertebrates and the latter infecting invertebrates including insects and crustaceans (Chinchar et al. 2017). In aquatic invertebrates, five iridoviruses have been described and have genomic data available: decapod iridescent virus 1 (DIV1) (Chinchar et al. 2017; Qiu et al. 2017; Xu et al. 2016), carnivorous sponge-associated iridovirus (CaSpA-IV) (Canuti et al. 2022), Pentanymphon antarcticum iridovirus (Bojko et al. 2024), Daphnia iridescent virus (DIV-1) (Toenshoff et al. 2018), and sergestid iridovirus (SIV) (Tang et al. 2007). Decapod iridescent virus 1 has been shown to be a major threat to crustacean aquaculture, causing considerable economic losses due to mass mortalities. The virus is able to infect a wide range of freshwater and marine species including crayfish (*Cherax quadricarinatus* and *Procambarus clarkii*), penaeid shrimp (e.g. *Penaeus monodon* and *P. vannamei*), freshwater prawns (e.g. *Macrobrachium rosenbergii*), and crabs (e.g. *Eriocheir sinensis*) (Reviewed in Liao et al. (2022)).

Pathologies associated with irido-like viruses have been reported in bivalves. Gill necrosis virus (GNV) was associated with gill necrosis in adult Portuguese Oyster, *Crassostrea angulata*, in France in the 1960s (Comps and Duthoit 1976), with similar aetiology also observed in the Pacific Oyster, *C. gigas* (Comps 1983). Irido-like virus particles (named haemocyte infection virus (HIV)) have also been associated with mortalities of *C. angulata* in France in the 1970s, in the absence of gill necrosis (Comps and Duthoit 1979). Irido-like viruses have also been shown to affect younger life stages of bivalves -Oyster velar virus disease (OVVD) has been associated with mortalities of larval *C. gigas* in hatcheries in North America in the 1970s and 1980s (Elston 1979; Elston and Wilkinson 1985). Despite the problems reported due to infection with irido-like viruses in the late 20^th^ century, they do not currently appear to be a threat to bivalve production, with few reports and no sequence data available for bivalve-infecting *Iridoviridae*.

Here we 1. characterise a novel iridovirus in cockles by high-throughput sequencing and genome annotation, 2. describe the virus phylogenetically in relation to other known *Iridoviridae*, 3. characterise infection with the novel iridovirus in its host by histopathology and transmission electron microscopy (TEM), and 4. identify the association between cockle moribundity and presence of the novel virus and *M. cocosarum*.

## 3 Methods

### 3.1 Sample collection

48 moribund cockles, characterised by an inability to burrow and a delayed reaction to stimuli, were collected from Dills Sand in the Wash Estuary, UK in May 2022 (Table 1). A small section of gill and mantle was dissected from each animal for molecular determination of infection with *M. cocosarum*, and the remaining tissues were maintained at 4°C prior to gradient purification of *Marteilia* cells. Samples for molecular, histopathology and electron microscopy analysis were collected in 2021 and 2023 as outlined in Table 1.

**Table 1:** Table outlining samples taken for this study. For “Sample type” column, M = Molecular, H = Histopathology, E = Electron microscopy, P = Purification of *Marteilia*.

| Sampling Site Name | Geographical location | Sample codes | Date(s) sampled | Moribund cockles (n) | Healthy cockles (n) | Sample type (M/H/E/P) |
| --- | --- | --- | --- | --- | --- | --- |
| Dills Sand | 52 56'.150 N<br>00 07'.050 E | RA21014 (50-89, 140-189) | 29/04/2021 | 40 | 50 | M/H/E |
|  |  | DS2022 | 16/05/2022 | 48 | 0 | M/P |
|  |  | RA23035 | 19/06/2023 | 50 | 50 | M/H/E |
| East Breast | 52 49'.870 N<br>00 20'.800 E | RA23040 | 06/06/2023 | 29 | 53 | M/H/E |
| Horseshoe Point | 53 30'.250 N<br>00 05'.500 E | RA23036 | 07/06/2023 | 50 | 50 | M/H/E |
| Inner West Mark Knock | 52 50'.300 N<br>00 14'.250 E | RA21022 | 27/07/2021 | 50 | 50 | M/H/E |
|  |  | RA23038 | 05/06/2023 | 50 | 50 | M/H/E |
| Mare Tail | 52 54'.950 N<br>00 07'.200 E | RA21014 (1-49, 90-139) | 29/04/2021 | 49 | 50 | M/H/E |
|  |  | MT2022 | 19/06/2023 | 48 | 0 | M/P |
|  |  | RA23037 | 19/06/2023 | 50 | 50 | M/H/E |
| Wrangle | 52 58'.800 N<br>00 08'.350 E | RA23039 | 08/06/2023 | 50 | 50 | M/H/E |

### 3.2 Gradient purification

DNA was extracted from gill and mantle tissue biopsies from 2022 samples as in (Tidy et al. 2025). All DNA extractions were subjected to a PCR using the MartDBITSf1 (5’-CTCGTGGAGCGGGTCTACCG-3’) and MartDBITSr1 (5’-TATCACGCCGCTGAATGCTTTCG-3’) primer set (Kerr et al. 2018), using the reaction composition and conditions outlined in Tidy et al. (2025). Animals that had a bright band of the correct size by agarose gel electrophoresis were selected for gradient purification.

Using the *M. cocosarum* presence data from the PCR, tissues from 34 animals were progressed to *Marteilia* purification using the method outlined in Robledo et al. (1995). Briefly, tissues were homogenised using an Ultra-Turrax homogeniser in 0.2 µm filtered seawater with 1% Tween80 (FSWT) in a total volume of 30 ml. The homogenates were then successively sieved through 250 and 75 µm nylon meshes. The resulting suspensions were centrifuged at 2,500 x *g* for 30 minutes at 4°C. The pellet was resuspended in 6 ml FSWT and deposited on a discontinuous 5-30% *(w/w)* sucrose gradient. After centrifugation at 2,500 x *g* for 30 minutes at 4°C, the 10-15%, 15-20% and 20-25% interfaces were collected, mixed, and diluted *v/v* in FSWT, then centrifuged at 2,500 x *g* for 10 minutes at 4°C to eliminate sucrose. The pellet, thought to contain *Marteilia* sporont primordia, was then resuspended in 200 µl 0.2 µm filtered seawater (FSW) and stored at -20°C until further processing.

For DNA extraction from the purification, 100 µl of the cells in FSW were added to 800 µl Lifton’s buffer (Winnepenninckx et al. 1993) in a lysing matrix A tube (MP Biomedicals) and homogenised at 5 m/s for 1 minute in a FastPrep24 Homogeniser (MP Biomedicals). Homogenates were incubated at 55°C for at least 3 hours with 20 µl 10 mg/ml Proteinase K (Sigma Aldrich). 100 µl of this homogenate was then extracted on a Maxwell48 Instrument (Promega) using the RSC DNA tissue kit (Promega), eluting into 100 µl elution buffer (Promega).

### 3.3 Metagenomic sequencing

DNA extracted from the gradient purification was prepared for metagenomic sequencing using the Nextera XT library preparation kit (Illumina, San Diego, CA, USA) following the manufacturer’s instructions, but using half volume reactions, and sequenced on an Illumina MiSeq using v3 chemistry and 2 × 300 bp cycles (Illumina).

### 3.4 Bioinformatic analysis

Raw Illumina paired-end sequence reads were trimmed to remove adaptor and low quality sequences using Trimmomatic v0.39 (in paired-end mode using a sliding window of 4, minimum quality of 15, leading and trailing values of 3, and a minimum length of 100 bases; Bolger et al. (2014)). The quality of trimmed and filtered reads was assessed using FastQC v0.11.9 (default parameters; https://www.bioinformatics.babraham.ac.uk/projects/fastqc/) prior to host removal using Bowtie2 v2.4.4 (Langmead and Salzberg 2012) and *C. edule* genome accession number GCA_947846245.1. Host-removed reads were assembled using SPAdes v3.15.3 (in –meta mode, using kmer sizes of 21, 33, 55, 77, 88 and 127; Prjibelski et al. (2020)). Assembled contigs were subsequently annotated using the BLASTp algorithm of Diamond (Buchfink et al. 2015) and the full NCBI non-redundant (nr) protein database (downloaded August 2021) and the results were visualised using MEGAN6 Community Edition v6.21.7. Paired reads were mapped to an assembled contig considered to represent a whole viral genome sequence of a novel iridovirus using BWA-MEM v0.7.17 and SAMtools v1.9 with default parameters (Li and Durbin 2009; Li et al. 2009). The output from mapping was visualised with Integrative Genomics Viewer (IGV) v2.5.2 (Robinson et al. 2011). Assembly quality and accuracy was assessed with QualiMap v2.2.2 (García-Alcalde et al. 2012).

Putative open reading frames (ORFs) were identified using four different tools: GeneMarkS using virus sequence type (Besemer et al. 2001), FGenesV (SoftBerry), Vgas (Zhang et al. 2019), and Prokka v1.14.0 in viral annotation mode (Seemann 2014). ORFs that were supported by two or more programmes were analysed further. Supported ORFs were annotated using NCBI BLASTp and the full NCBI protein sequence database (accessed August 2023), OrthoFinder and a representative subset of iridovirus proteins (using Diamond for sequence searching and MAFFT for sequence alignment; Emms and Kelly (2019)), and protein motifs were identified by HHpred (default parameters) and InterProScan 5.

### 3.5 Phylogenetic analysis

Six core iridovirus genes identified by Toenshoff et al. (2018) were used for phylogenetic analysis: the major capsid protein (BiIV1_119R), DNA-directed RNA polymerase II subunit RPB2 (BiIV1_105L), a putative A32-like packaging ATPase (BiIV1_063R), a putative CTD phosphatase-like protein (BiIV1_153L), a putative helicase protein (BiIV1_025R), and a putative transcription elongation factor S-II-like protein (BiIV1_108L). Proteins from a representative set of 114 *Iridoviridae* (including BiIV1) and five *Ascoviridae* were used for the analysis, with two members of *Marseilleviridae* used as an outgroup. The whole viral genomes from which the proteins originated are outlined in Supplementary Table S1. Orthologues of the six core genes were identified and aligned with OrthoFinder (parameters as above; Emms and Kelly (2019)). Resulting alignments were checked by eye and concatenated into a single alignment for further analysis. A Bayesian consensus tree was constructed from the concatenated protein alignment using MrBayes v3.2.7 (Ronquist et al. 2012) on the CIPRES Science Gateway (Miller et al. 2010). The tree was constructed using two separate MC^3^ runs, carried out for two million generations using one cold and three hot chains. The first 500,000 generations were discarded as burn-in, and trees were sampled every 1000 generations.

### 3.6 PCR

DNA was extracted from tissues (a section comprising gill, mantle and digestive gland) fixed in RNAlater (Sigma Aldrich) from 2021 and 2023 samples, as in Tidy et al. (2025), for PCR screens for BiIV1 and *Marteilia*. Nested PCR primers were designed to amplify 573 bp and 462 bp regions of the ATPase gene of BiIV1 (Table 2). PCR reactions were carried out in 20 µl reactions containing of 1 μl DNA template, 1× Green GoTaq® Flexi Buffer (Promega, Wisconsin, USA), 2.5 mM MgCl_2_ (Promega), 0.4 mM dNTP mix (Bioline, London, UK), 0.5 µM forward primer, 0.5 µM reverse primer, 10 µg Bovine Serum Albumin (BSA) (New England Biolabs, Massachusetts, USA), and 0.5 u GoTaq® G2 DNA Polymerase (Promega). The first round PCR product was amplified using the using BiIV1_ATPase_f1/BiIV1_ATPase_r1 primer set, by an initial denaturing at 95°C for 5 minutes, followed by 35 cycles of 95°C for 45 seconds, 60°C for 45 seconds, and 72°C for 45 seconds; and a final extension of 10 minutes at 72°C. The nested amplicon was produced using the BiIV1_ATPase_f2/BiIV1_ATPase_r2 primer set with cycling conditions as above, but with an annealing temperature of 58°C. A Marteilia-specific nested PCR, targeting the ITS1 region, was performed with the same reaction components as above, using the nested primer set (MartDBITSf1/MartDBITSr1 and MartDBITSf2/MartDBITSr2) and cycling conditions of Kerr et al. (2018).

**Table 2:**
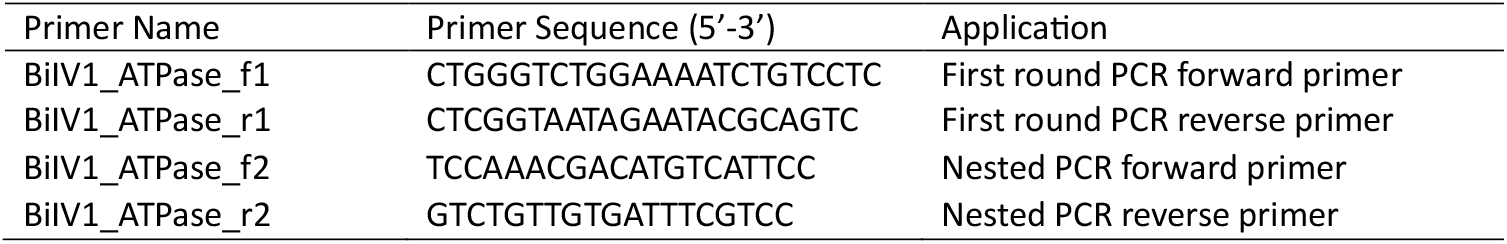
Primer sequences used to amplify the ATPase gene of BiIV1.

### 3.7 Histopathology and transmission electron microscopy

Samples fixed for histopathology were collected in 2021 and 2023 (Table 1). Tissue cross-sections of whole animals were fixed in Davidson’s seawater fixative for 24-48 hours before transferring to 70% industrial denatured alcohol (IDA). Fixed tissues were processed routinely for histology, sectioned, and stained with haematoxylin and eosin (H&E). Sections were examined using a Nikon Eclipse E800 light microscope and digital images captured using NIS imaging software.

For TEM, small pieces of gill and mantle tissue (*ca*. 2 mm^3^) were fixed in 2.5% glutaraldehyde in 0.1 M sodium cacodylate buffer (pH 7.2). Post-fixation was carried out in 1% osmium tetroxide/0.1 M sodium cacodylate buffer for one hour, followed by washing twice with 0.1 M sodium cacodylate buffer. Fixed tissues were dehydrated through a graded acetone series prior to embedding in Agar 100 epoxy resin (Agar Scientific). Semi-thin (1–2 µm) sections were stained with Toluidine blue for viewing with a light microscope to identify sections with suitable target regions. Ultra-thin sections (70–90 µM) of target regions of the tissues were mounted on uncoated copper grids and stained with 2% aqueous uranyl acetate and Reynold’s lead citrate (Reynolds 1963). Grids were examined using a JEOL JEM1400 transmission electron microscope and digital images captured using an AMT XR80 camera and AMT V602 software.

## 4 Results

### 4.1 Sequence analysis

#### 4.1.1 Complete Genome Assembly of BiIV1

A total of 19,541,012 Illumina read pairs were generated from the library; after quality-trimming and filtering 19,185,146 forward and 18,026,547 reverse reads remained. Assembly of trimmed and filtered reads using SPAdes produced a 179,695 bp contig with similarity to *Iridoviridae* using Diamond

Blastx analysis. We henceforth refer to this novel iridovirus as bivalve iridovirus 1 (BiIV1). Mapping trimmed and quality filtered reads back to the consensus BiIV1 genome showed an average coverage of 1,209×. The full genome was deposited to NCBI GenBank under accession number PQ846775.

Gene-prediction software suggested that BiIV1 contains 193 open reading frames (ORFs), which were supported by ≥2 prediction programmes, with 90 on the forward stand and 103 on the reverse strand (Figure 1). The G+C content of BiIV1 was calculated to be 40.99%, similar to other *Betairidiovirinae*, which are typically characterised as having G+C content lower than 50% (Chinchar et al. 2017). The genome size of BiIV1 was also similar to that of other *Betairidovirinae*. A comparison of the G+C content and genome size of BiIV1 to representatives of *Alpha- and Beta-iridovirinae* is shown in Figure 2.

**Figure 1.**
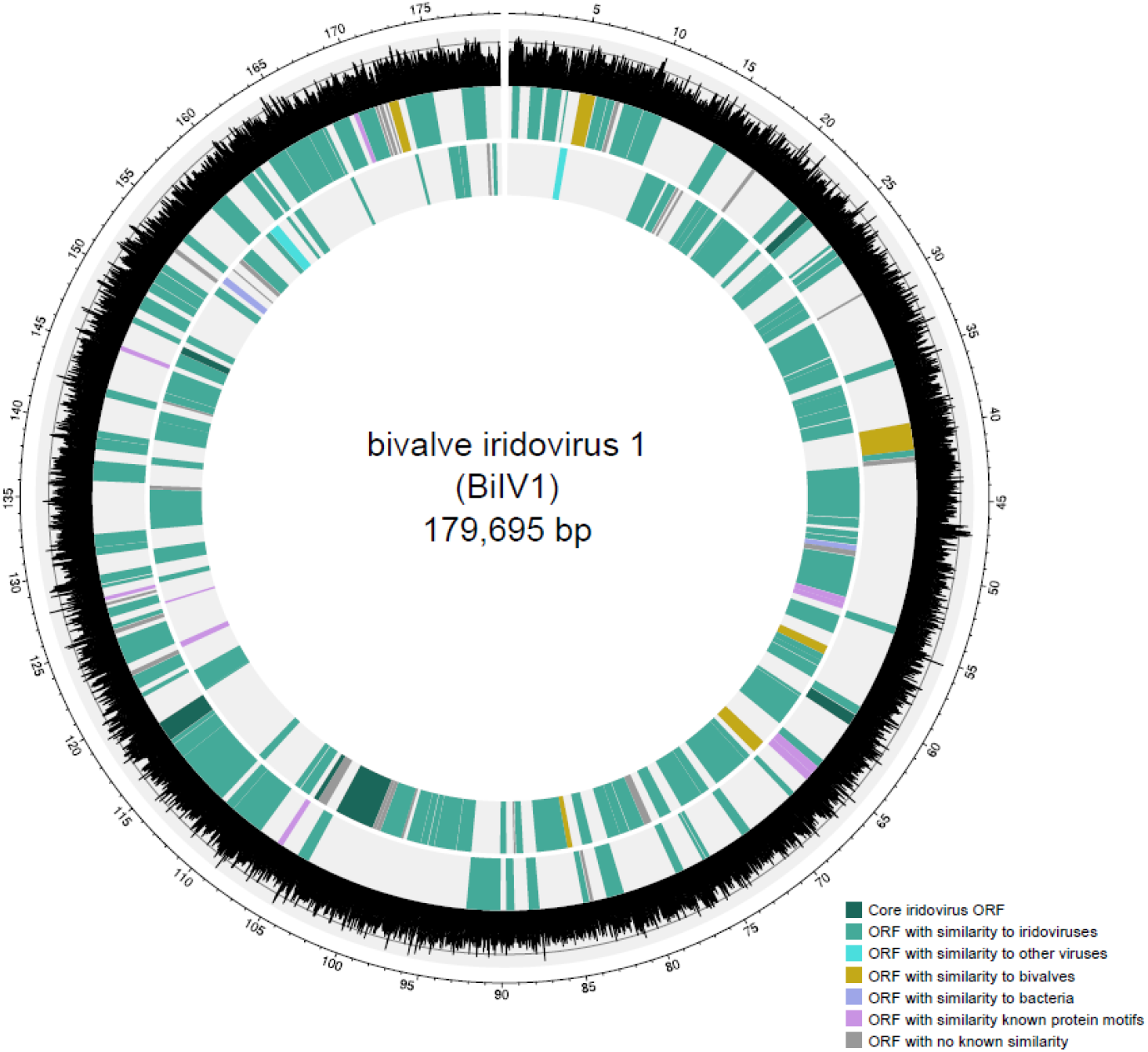
Circular map of the 179,695 bp bivalve iridovirus 1 (BiIV1) genome. Outer scale is numbered clockwise in kb. Outer graph in black and grey depicts G+C content (%) across the genome. The second innermost track depicts predicted open reading frames (ORFs) on the positive strand, and the innermost track depicts the predicted ORFs on the negative strand. Dark green blocks represent ORFs with homology to the six core genes identified by Toenshoff et al. (2018), mid-green blocks represent ORFs with similarity to other iridoviruses, light blue blocks represent ORFs with similarity to other viruses and yellow blocks represent ORFs with predicted protein sequence similar to bivalve proteins. Purple blocks represent ORFs with similarity to bacterial-derived proteins, pink blocks depict ORFs that contained sequences consistent with known protein motifs, and ORFs represented by grey blocks had no similarity to known proteins or protein motifs.

**Figure 2.**
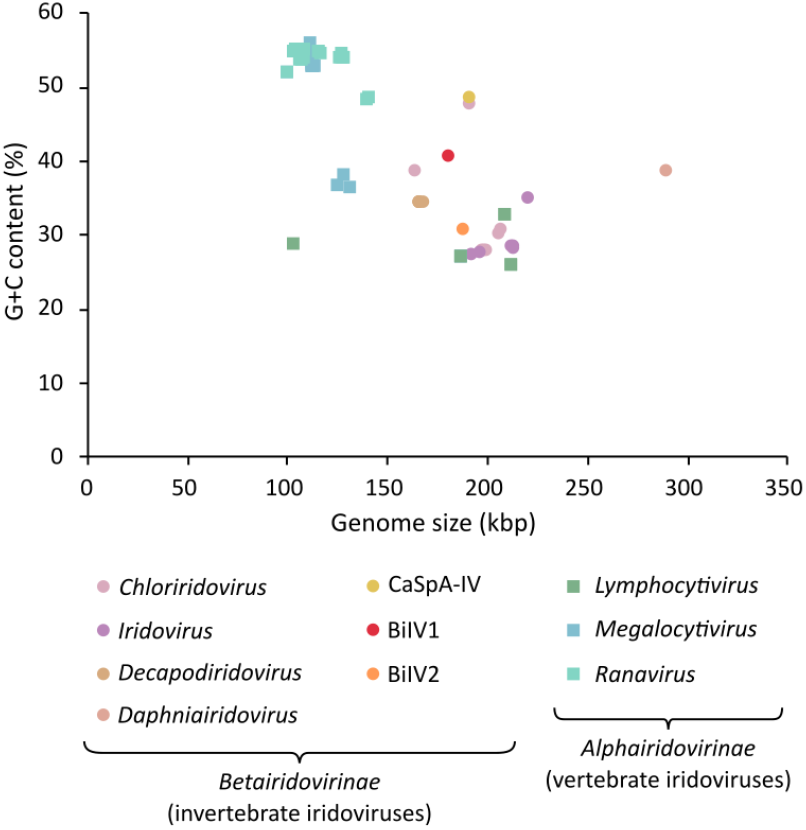
Genome size (kbp) plotted against G+C content (%) for a representative set of *Iridoviridae* genomes. *Alphairidovirinae* are represented by squares, with genera within this subfamily represented by different colours. *Betairidovirinae* are represented by circles, with the four defined genera within this subfamily, and the three species within no assigned genus (including BiIV1), represented by different colours. Within the *Betairidovirinae* subfamily, Anopheles minimus iridovirus (accession number KF938901) has the smallest genome size at 163,023 kbp, and Daphnia iridovirus 1 (accession number LS484712) has the largest at 288,858 kbp. G+C content within *Betairidovirinae* ranges between 27.75% (cricket iridovirus – accession number OK181107) and 48.75% (carnivorous sponge associated iridovirus – accession number ON887238).

As part of this study, the major capsid protein of BiIV1 was used to search the whole-genome shotgun (wgs) database for similar sequences in bivalve and crustacean datasets. This revealed a 187,132 bp contig (accession number CAUJNT010007968), suspected to be another novel iridovirus, from a sequencing dataset of a deep-sea mussel, *Bathymodiolus septemdierum*. We henceforth refer to this putative viral genome as bivalve iridovirus 2 (BiIV2) and use its genome and predicted protein sequences to compare to BiIV1.

#### 4.1.2 Genome features of BiIV1

##### 4.1.2.1 Annotation of the BiIV1 open reading frames

The 193 predicted ORFs were annotated by comparison to NCBI nr, InterproScan and UniProtKB databases, and by orthologue analysis. 151 ORFs showed similarity, either by blastx or homologue analysis, to iridovirus proteins, two ORFs had similarly to other virus families, two had similarity to bivalve proteins, and two had similarity to bacterial proteins. Three of the ORFs that had similarity to iridovirus proteins also had similarity to bivalve proteins. Of the remaining 41 ORFs, 26 had no predicted protein motif similarity by analysis with InterproScan or by blasting against the UniProKB database.

##### 4.1.2.2 BiIV1 core iridovirus genes

Orthologue analysis, carried out by comparing the 193 predicted BiIV1 genes to the predicted genes of 113 other *Iridoviridae*, suggested that 20 out of the 23 core genes proposed by Toenshoff et al. (2018) were present in the BiIV1 genome. Absent from the BiIV1 genome were two myristoylated membrane proteins, and a protein kinase. The same three core genes were also absent from the genome of BiIV2. All six genes that Toenshoff et al. (2018) suggested are present in all iridoviruses are present in both BiIV1 and BiIV2 and were used for phylogenetic analysis.

##### 4.1.2.3 Gene homology to other iridoviruses

Of the 193 predicted BiIV1 ORFs, 151 showed similarity, either by blastx or homologue analysis, to iridovirus proteins. Of these, 124 could be attributed to a function or had protein motif similarity by analysis against InterproScan or UniProtKB databases (Supplementary Table S2). Orthologue analysis revealed that of the 102 BiIV1 ORFs that had iridovirus orthologues, and all bar one shared protein similarity to *Betairidovirinae* or *Betairidovirinae* proteins that also have orthologues in *Alphairidovirinae*. The single ORF with an orthologue only present in *Alphairidovirinae*, BiIV1_084R, is not annotated, and has no matches to known protein motifs. 49 of the predicted iridovirus orthologues appear to be specific to aquatic invertebrate-associated iridoviruses, with these predicted genes absent from terrestrial *Betairidovirinae* and *Alphairidovirinae*. These ORFs included a predicted glycosyltransferase, a cytosine-specific methyltransferase, and a putative HD domain-containing protein. Of the 49 ORFs that only have orthologues in aquatic invertebrate-associated iridoviruses, 17 appear to be only present in BiIV1 and BiIV2, suggesting that they may be specific to bivalve-associated iridoviruses. Alternatively, some of these 17 ORFs may be too distantly related to other iridovirus proteins to be classified as an orthologue; for example, BiIV1 and BiIV2 predicted DNA-directed RNA polymerase subunit, Rpb5, was not binned with other *Betairidovirinae* Rpb5 ORFs.

##### 4.1.2.4 Candidates for horizonal gene transfer between BiIV1 and its host

As *Iridoviridae* are known to possess genes that encode proteins orthologous to their hosts, acquired through horizonal gene transfer (HGT) between the virus and its host and/or ancestral hosts (Becker and Darai 2012), we further investigated whether the predicted ORFs of BiIV1 possess homology to bivalve genes to determine if they could have been acquired by HGT. Nine BiIV1 ORFs had homology to *C. edule* or other bivalve proteins (Table 3). The ORF with the greatest homology to bivalve proteins was BiIV1_058L, with 79.7% sequence similarity across the predicted coding sequence to its cockle homologue. BiIV1_058L is predicted to encode a homologue of charged multivesicular body protein 4 (CHMP4). BiIV1_044R and BiIV1_069L are predicted to contain domains, similar to EGF domains in *Mytilus* species. Two BiIV1 ORFs have similarity to bivalve proteins involved in DNA synthesis and replication: BiIV1_087R is predicted to encode dihydrofolate reductase, involved in DNA precursor biosynthesis, and BiIV1_090R is predicted to encode a proliferating cell nuclear antigen. BiIV1_006R and BiIV1_186R are predicted to be homologues of bivalve proteins in involved in the regulation of apoptosis and cell-cycle. BiIV1_006R is predicted to encode a protein with a caspase recruitment domain (CARD) and RIP homotypic interaction motif (RHIM) domain with similarity to bivalve-derived sequences, and BiIV1_186R is predicted to encode a cyclin-like protein, with similarities in protein sequence to bivalve G1/S cyclins. Two further BiIV1 predicted proteins, BiIV1_039R and BiIV1_088L, had similarly to bivalve proteins, however the function of these proteins is currently uncharacterised. Of the nine BiIV1 genes that had similarity to bivalve proteins, six of them had BiIV2 homologues: BiIV1_006R, BiIV1_039R, BiIV1_044R, BiIV1_069L, BiIV1_090R, and BiIV1_186R.

**Table 3:**
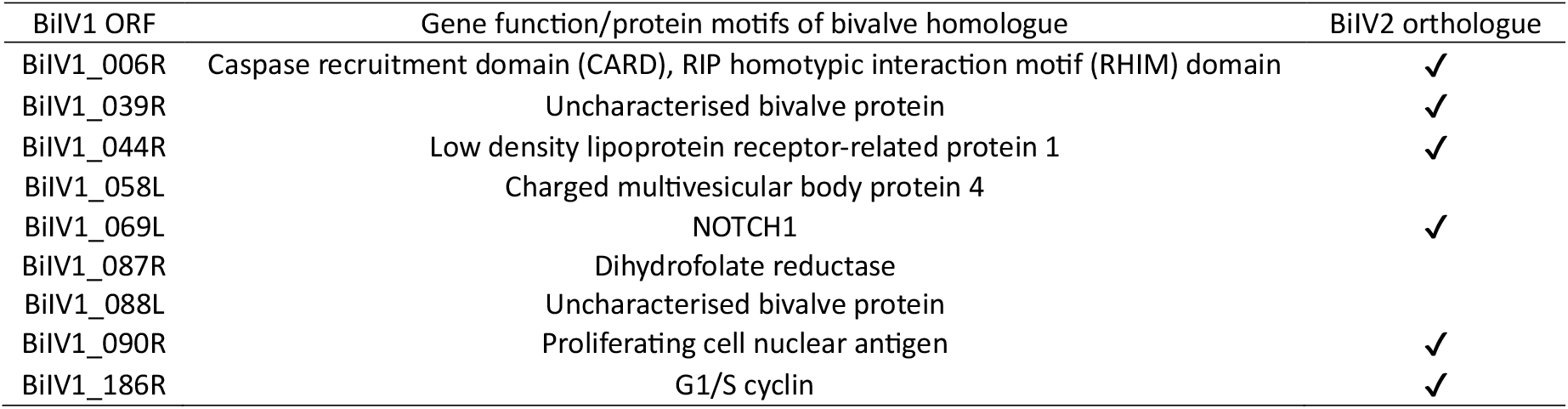
Candidate genes for horizonal gene transfer between *Cerastoderma edule*, its ancestors, and bivalve iridovirus (BiIV1). Presence of orthologues of the candidate genes in a second putative iridovirus, bivalve iridovirus 2 (BiIV2) is also presented.

#### 4.1.3 Phylogenetic Analysis

To compare the phylogenetic relationship of BiIV1 to other members of *Iridoviridae*, the six core genes from BiIV1 were aligned against a subset of *Iridoviridae*, representing all known iridoviruses (where full genomes exist) and their hosts, as well as a representative set of *Ascoviridae*. BiIV1 and BiIV2 branched as a maximally-supported sister clade to that containing most other aquatic invertebrate iridoviruses: the *Decapodiridovirus* genus, carnivorous sponge-associated iridovirus (CaSpA-IV), and Pentanymphon antarcticum iridovirus (Figure 3). The only other iridovirus known to infect aquatic invertebrate hosts, Daphnia iridescent virus 1 (DIV-1), did not branch closely to all other aquatic iridoviruses and was a sister clade to the *Chloriridovirus* genus, comprising iridoviruses with semi-aquatic hosts.

**Figure 3.**
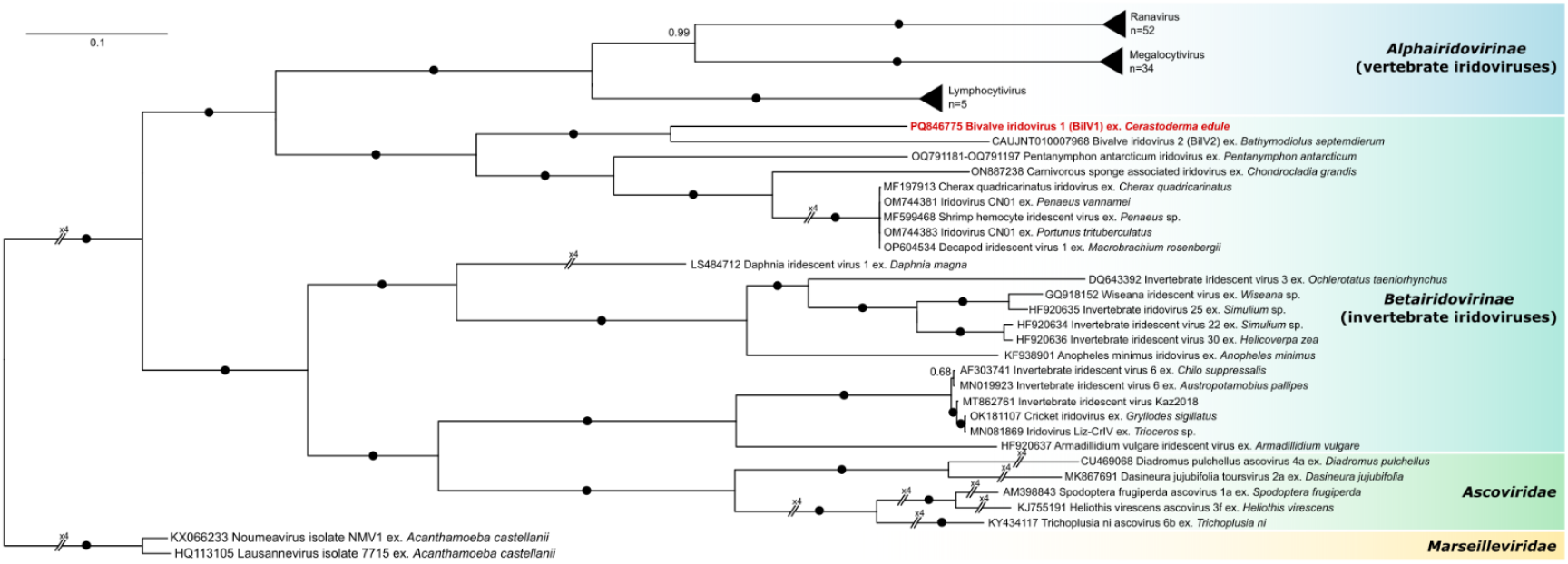
Bayesian consensus tree constructed from a concatenated multiple amino acid alignment of major capsid protein, DNA-directed RNA polymerase II subunit Rpb2, putative A32-like packaging ATPase, putative CTD phosphatase-like protein, putative helicase protein, and putative transcription elongation factor S-II-like protein from BiIV1, 113 other *Iridoviridae*, five *Ascoviridae* and two *Marseilleviridae*. The tree is rooted to the *Marseilleviridae* clade. Branch labels denote posterior probabilities, with black circles used when posterior probability = 1.

#### 4.1.4 Transmission electron microscopy and histopathology

Icosahedral virions were observed in the cytoplasm of infected haemocyte cells, often in areas directly adjacent to cell nuclei which displayed condensed chromatin. Fully formed virions possessed a clear capsid, electron dense core and an intermediate amorphous layer containing an internal lipid membrane. The mean particle sizes were 165 nm vertex to vertex, 151 nm face to face, and had an average equivalent diameter of 158 nm (n=50). Virion assembly occurs within the cytoplasm of infected cells, developing and empty capsids were observed alongside fully formed virions, and accumulations of putative viral DNA (Figure 4).

**Figure 4.**
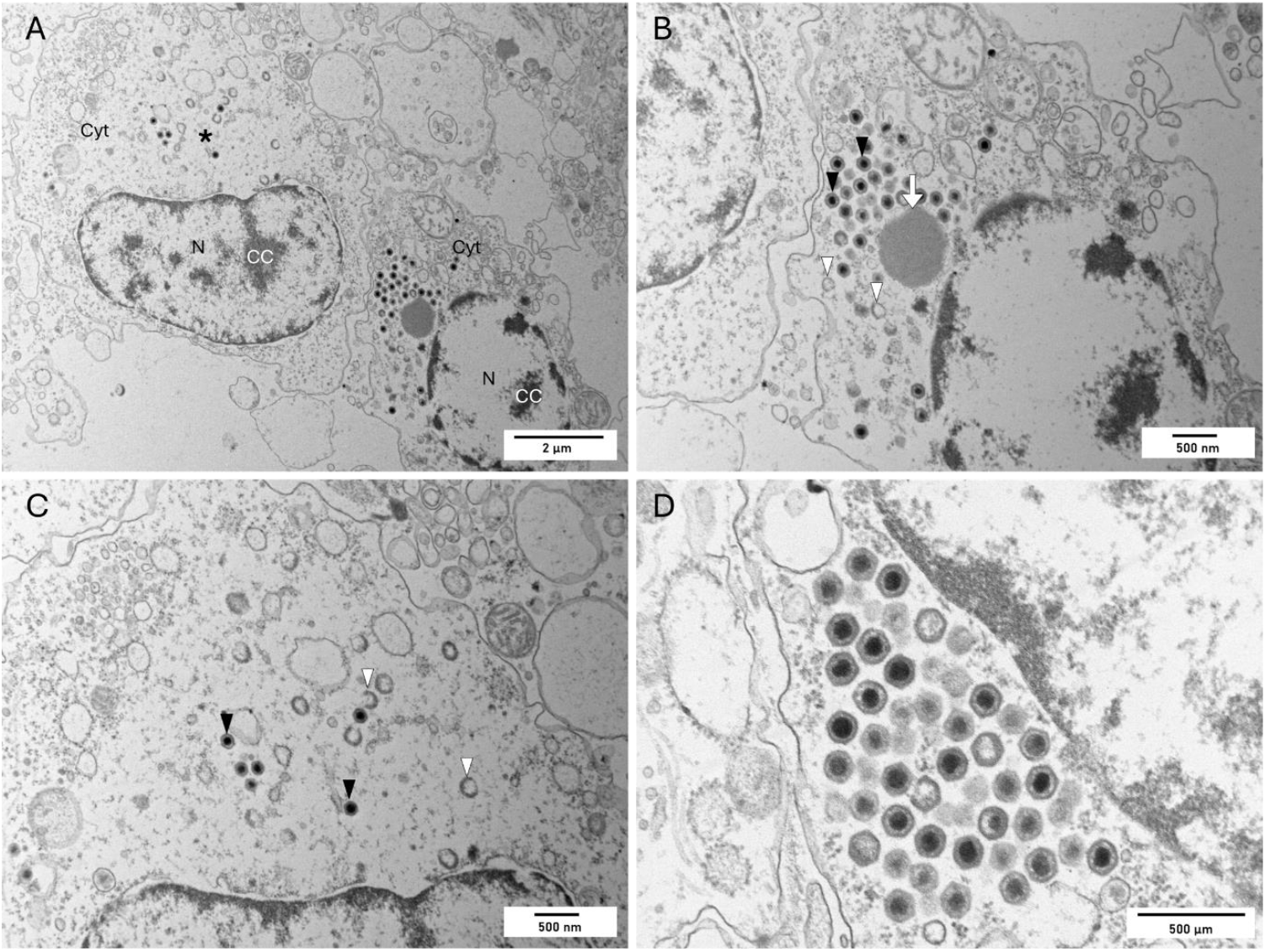
Electron micrographs of a common cockle, *Cerastoderma edule*, infected with bivalve iridovirus 1 (BiIV1). (A) Two host cells infected with BiIV1, with the nuclei, N, showing condensed chromatin (CC), and mature and immature BiIV1 virions in the cytoplasm (Cyt). The left cell has a large electron-lucent assembly site (*) containing a few mature virions and many immature, developing virions. The right cell contains a paracrystalline array of virus particles. Scale bar = 2 µm. (B) Higher magnification of the right cell, with numerous mature virions (black arrowheads), a few immature, developing virions (white arrowheads), and putative viral DNA prior to packing (white arrow). Scale bar = 500 nm. (C) Higher magnification of the left cell, with mature virions (black arrowheads) and immature, empty and developing capsids (white arrowheads). Scale bar = 500 nm. (D) High magnification of mature virions in a paracrystalline array, showing detail of the envelope, intermediate amorphous layer, and electron dense core of BiIV1. Scale bar = 500 nm.

Histopathology highlighted haemocytic infiltration within the connective tissues surrounding the digestive gland tubules. Some of these cells were observed to contain basophilic inclusion bodies within the cytoplasm, pathology similar to that described for DIV1 infections in shrimp tissues. Nuclei of the affected cells were observed to possess condensed and marginalised chromatin (Figure 5). As highlighted by Tidy et al. (2025), inflammation characterised by haemocytic infiltration was noted in 78% of moribund cockles and 24% of healthy (buried cockles) sampled at Mare Tail and 42% of healthy cockles at Dills Sand. Haemocytic infiltration was noted to be associated with presence of *Marteilia*-like cells in 60% of the moribund cockles, 24% of the healthy cockles at Mare Tail and 12% of the healthy cockles at Dills Sand. Interestingly it was the animals which displayed a clear host response without the presence of *Marteilia*-like cells where we observed viral infection.

**Figure 5.**
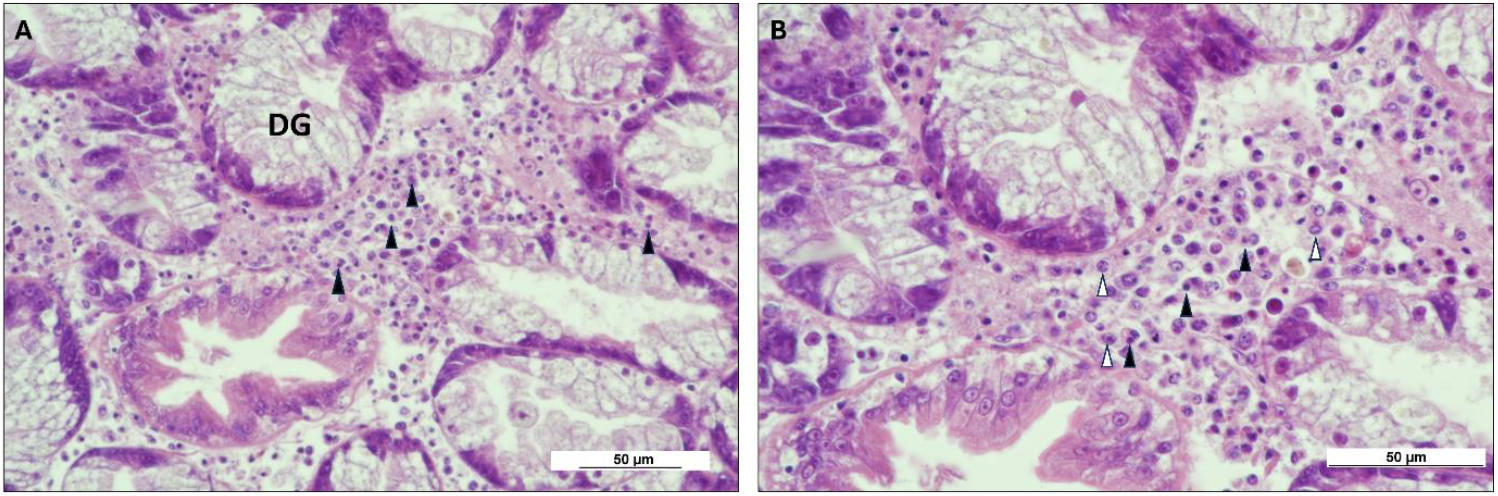
Histopathology images of *Cerastoderma edule* infected with bivalve iridovirus 1 (BiIV1). A) Infiltrating haemocytes within the connective tissues surrounding the digestive gland tubules (DG) were observed to possess basophillic inclusions (arrowheads) in the cytoplasm. Scale bar = 50 µm. B) Higher magnification of haemocytes with basophilic inclusions (black arrowheads) within the cytoplasm. Nuclei of the affected cells also showed marginated and condensed chromatin (white arrowheads). Scale bar = 50 µm.

### 4.2 PCR Prevalence of BiIV1 and Marteilia

The prevalence of BiIV1 and *Marteilia* was assessed by PCR. For all sampling events except Dills Sand in 2023, the prevalence of BiIV1 by PCR was higher in moribund cockles than in apparently healthy cockles. Two sites in 2023 appeared to have a very high occurrence of BiIV1: East Breast and Horseshoe Point had BiIV1 prevalences of 100% and 90% in moribund cockles, and 76.5% and 58% in apparently healthy cockles, respectively (Table 4). Strong positives from the first round of PCR were also seen to be more prevalent in moribund cockles from most sites (data not shown). Some sampling points appeared to have a lower overall presence of BiIV1 than others, for example, Mare Tail in 2023 had a 14% PCR prevalence of BiIV1 in moribund cockles and a 4% prevalence in apparently healthy cockles; this is in contrast to the previous two sampling years where PCR presence in moribund cockles was 78% and 96% in 2022 and 2021, respectively.

**Table 4.** PCR prevalence of bivalve iridovirus 1 (BiIV1) and *Marteilia cocosarum* in moribund and apparently healthy cockles sampled in 2021-2023. PCR prevalence of *Marteilia* from Dills Sand, Inner West Mark Knock and Mare Tail in 2021 were determined in Tidy et al. (2025).

| Site Name | Date sampled | Moribund cockles (n) | Healthy cockles (n) | BiIV1 PCR prevalence in moribund cockles (%) | BiIV1 PCR prevalence in healthy cockles (%) | *Marteilia* PCR prevalence in moribund cockles (%) | *Marteilia* PCR prevalence in healthy cockles (%) | Co-occurrence of BiIV1 and *M. cocosarum* (%) |
| --- | --- | --- | --- | --- | --- | --- | --- | --- |
| Dills Sand | 29/04/2021 | 40 | 50 | 70 | 10 | 95 | 42 | 44.4 |
|  | 16/05/2022 | 48 | 0 | 96 | - | 76 | - | 58.33 |
|  | 19/06/2023 | 50 | 50 | 2 | 10 | 4 | 8 | 0 |
| East Breast | 06/06/2023 | 29 | 51 | 100 | 76.5 | 0 | 13.7 | 6.25 |
| Horseshoe Point | 07/06/2023 | 50 | 50 | 90 | 58 | 82 | 48 | 35 |
| Inner West Mark Knock | 27/07/2021 | 50 | 50 | 6 | 0 | 22 | 10 | 0 |
|  | 05/06/2023 | 50 | 50 | 28 | 20 | 42 | 12 | 9 |
| Mare Tail | 29/04/2021 | 49 | 50 | 78 | 20 | 83.7 | 34 | 49.5 |
|  | 16/05/2022 | 48 | 0 | 96 | - | 39.6 | - | 35.4 |
|  | 19/06/2023 | 50 | 50 | 14 | 4 | 30 | 24 | 8 |
| Wrangle | 08/06/2023 | 50 | 50 | 72 | 22 | 0 | 0 | 0 |

When compared to the prevalence of *Marteilia*, BiIV1 appeared to occur at a similar frequency as *Marteilia* at Dills Sand, Horseshoe Point, Inner West Mark Knock, and most sampling years at Mare Tail sites (i.e. when BiIV1 was abundant so was *Marteilia*, and when BiIV1 appears at low levels, so does *Marteilia*). However, this pattern was not observed at the Wrangle site in 2023: 72% and 22% PCR prevalence of BiIV1 in moribund and healthy cockles, respectively; and at East Breast: 100% prevalence of BiIV1 and an absence of *Marteilia* was observed in moribund cockles in 2023. To a lesser extent, this was also observed at the Mare Tail site in 2023, with a 96% PCR prevalence of BiIV1 in moribund cockles, versus a 39.6% presence of *Marteilia*. Co-presence of BiIV1 and *Marteilia* was common where both were present at a site, for example, at the Mare Tail site in 2021, 32/49 moribund cockles were PCR positive for both BiIV1 and *Marteilia*, but 5 animals were PCR positive for BiIV1 only, and 9 animals were PCR positive for *Marteilia* only.

## 5 Discussion

In this study, we present a novel member of the *Iridovirdae* family, bivalve iridovirus 1 (BiIV1), characterising its genome, morphology, and presence in host cells. BiIV1 represents the third iridovirus infecting an aquatic invertebrate to be characterised by genomics and histopathology/TEM, with genomes and corresponding evidence of infection available only for decapod iridescent virus 1 (DIV1) and daphnia iridescent virus (DIV-1). The data produced in this study adds to the knowledge of iridovirus genomics and diversity, particularly those associated with aquatic invertebrates.

Despite reports of pathologies associated with irido-like viruses in the *Crassostrea* genus of bivalves in the 1970s and 80s (Comps 1983; Comps and Duthoit 1976; 1979; Elston 1979; Elston and Wilkinson 1985), no genomic data for these exist. In this study, we produced a genome for BiIV1, but also discovered a second bivalve-associated iridovirus, coined bivalve iridovirus 2 (BiIV2), in whole genome sequencing data for *B. septemdierum*, a species of deep sea mussel (Hashimoto and Okutani 1994). *B. septemdierum* is an important foundation species that form high biomass beds in hydrothermal vent ecosystems in the western Pacific and Indian Oceans (Tunnicliffe and Breusing 2022). One chromosome-level genome and one alternative haplotype dataset exist for *B. septemdierum*. The alternative haplotype is a contig-level assembly, comprising 8,524 scaffolds, one of which, accession number CAUJNT010007968, we propose to be BiIV2. The chromosome level assembly has 129 unplaced scaffolds, one of which, accession number CAUJNP010000008, we believe to contain *ca*. 3.3 copies of the BiIV1 genome, potentially misassembled due to the circularly permuted nature of *Iridoviridae* genomes (Chinchar et al. 2017).

BiIV1 has similarities in genome size and protein sequence, virion morphology and mode of infection to other *Iridoviridae*. 78.2% of the 193 predicted BiIV1 ORFs shared similarities in protein sequence to other *Iridoviridae* by either Blastx search, protein motif similarity, or using orthologue analysis. By orthologue analysis, BiIV1 exclusively shared protein similarity to *Betairidoririnae* or *Betairidovirinae* proteins that also have homologues in *Alphairidovirinae*, bar one ORF. BiIV1 appeared to contain 17 genes that are exclusive to bivalve iridoviruses, including six genes that are potentially homologues acquired from their hosts. Virion morphology of BiIV1 is typical of that of other *Iridoviridae*, with BiIV1 virions approximately 158 nm in diameter. This virion size is very similar to that of DIV1 (Arulmoorthy et al. (2022)), but much smaller than the virions previously reported to infect other species of bivalve, which ranged from 288 – 380 nm (Comps and Duthoit 1976; 1979; Elston 1979). BiIV1 showed similarities to other *Iridoviridae* in regard to virion assembly: Potential virus-derived DNA and virion assembly observed in the cytoplasm of the host haemocytes, with similar pathology to that of DIV1 (Xu et al. 2016). Mature virions were also seen free in the hemocoel, suggesting that BiIV1 is budding from the haemocytes, although this was not observed by TEM.

A number of viruses, including *Iridoviridae*, are known to possess genes that encode proteins homologues to their host and/or its ancestors, acquired through horizonal gene transfer (HGT) (Becker and Darai 2012). Publicly available genomes for *C. edule* are currently unannotated, therefore orthologue analysis could not be carried out between BiIV1 and its host. Instead, BiIV1 predicted protein sequences were searched against bivalve non-redundant databases, and any BiIV1 proteins with motifs similar to bivalve protein-coding sequences were investigated. All BiIV1 proteins predicted to have been acquired by horizonal gene transfer were intronless, but had similarity to the exon regions of bivalve genes, suggesting that they had been acquired by retrotransposition i.e. the reverse transcription of host mRNA (Vallée et al. 2021). BiIV1 ORFs thought to have been acquired by this mechanism are predicted to encode a range of functions, including those related to endosomal sorting complexes, DNA synthesis and replication, and cell cycle control. We discuss some of the genes in more detail below.

BiIV1_058L is predicted to encode a charged multivesicular body protein 4 (CHMP4). CHMP4 is a member of the Snf7 family of proteins, which forms part of the endosomal sorting complexes required for transport (ESCRT). Other enveloped viruses, such as human immunodeficiency virus (HIV), rely on certain components of the ESCRT machinery for viral budding – with only CHMP4 and CHMP2 appearing to be essential for this process (Schmidt and Teis 2012). The homologue of *C. edule* CHMP4 in BiIV1 may be able to exploit the host ESCRT machinery to bud out of the infected host cell. BiIV1_186R is predicted to encode a cyclin-like protein. The predicted protein sequence showed blast similarity to bivalve G1/S cyclins. Homologues of D-type cyclins, which function in the G1/S stage of cell proliferation, have been identified in gammaherpesviruses. These homologues have been shown to share functions with the host’s cellular cyclin, but some viral cyclins have also acquired altered characteristics (Hoge Ann et al. 2000). BiIV1_006R is predicted to encode a protein with a caspase recruitment domain (CARD) and RIP homotypic interaction motif (RHIM) domain with similarity to bivalve-derived sequences. Other iridoviruses have been found to possess CARD domains (Ke and Zhang 2023), and RHIM domains are present in herpesviruses (Baker et al. 2020); both domains have been shown to aid in inhibiting activation of host cell death pathways (Baker et al. 2020; Ke and Zhang 2023). BiIV1_006R and BiIV1_186R have orthologues in BiIV2, but are absent from all other known iridoviruses, suggesting that these protein-coding genes may be specific to iridoviruses that are associated with bivalves.

A previous study associated cockle moribundity in the Wash Estuary to the presence of *M. cocosarum* and disseminated neoplasia (Tidy et al. 2025). This study provides evidence that BiIV1 may be another aetiology of the mortalities that have been occurring yearly in the Wash since the initial observation in 2008. Our new findings explain the cause of inflammation present in some cockles in the absence of the *Marteilia* parasite in Tidy et al. (2025). Co-occurrence of BiIV1 and *Marteilia* was common in cockles at sites where the two pathogens were present. However, the presence of the pathogens singularly was also observed by histology and PCR, demonstrating that they can infect separately, and can cause host responses independently of each other. Cockles from sites that were sampled over multiple years appear to have differing prevalences of BiIV1 and *Marteilia* over time. This could be explained by the difference in sampling time each year; further temporal sampling across a year may elucidate whether time of year influences the presence of either pathogen. Some sites appear to have a lower overall presence of BiIV1 and *Marteilia*. Currently, the cause of all mortalities at these sites remains uncertain, with the presence of disseminated neoplasia unable to fully explain the levels of cockle moribundity reported. Environmental factors which differ between sites may explain discrepancies between the difference in the presence of pathogens and cockle pathology between sites, however such multifactorial investigations were outside of scope of this study.

## 6 Summary

We provide evidence that a novel iridovirus, bivalve iridovirus 1 (BiIV1), is associated with mortality events in cockles in the Wash Estuary, UK. Previously, mortalities had been attributed, at least in part, to infection with *Marteilia cocosarum*, and the presence of disseminated neoplasia, however their presence could not be attributed to all pathology observed in moribund cockles. We provide the full genome for BiIV1 and characterise its infection in *Cerastoderma edule* by electron microscopy and histopathology. We also identify a second bivalve-associated iridovirus (BiIV2) in whole genome sequence data from the deep-sea mussel, *Bathymodiolus septemdierum*. BiIV1 and BiIV2 are the currently the only bivalve-associated viruses with genomic sequence data available – they appear to contain genes specific to bivalve-associated iridoviruses, and also contain genes that are predicted to have been obtained through horizontal gene transfer between their hosts and host’s ancestors. This study enhances our knowledge of the diversity of invertebrate iridoviruses as well as providing further evidence that the mortalities occurring in the Wash are due to multiple stressors, and that a broad approach is required to understand how these stressors interact to result in bivalve mortalities.

## 7 Data availability

The full, annotated genome of bivalve iridovirus 1 (BiIV1) has been deposited to GenBank under accession number PQ846775.

## 8 Funding

This work was funded by Defra project FB002 and Cefas Seedcorn project DP1000.

## Notes

### Competing Interest Statement

The authors have declared no competing interest.

